# Skeletal muscle mitochondrial responses to a single bout and six weeks of high load versus high volume resistance training in previously trained men

**DOI:** 10.1101/2025.07.23.666423

**Authors:** Breanna Mueller, Carlton D. Fox, Hailey A. Parry, Paulo H. C. Mesquita, Christopher G. Vann, Bradley A. Ruple, Casey L. Sexton, Joshua S. Godwin, Mason M. McIntosh, Darren T. Beck, Kaelin C. Young, Stuart M. Phillips, Andreas N. Kavazis, Michael D. Roberts

**Author notes:** Address co-correspondence to: Michael D. Roberts, PhD, Professor, School of Kinesiology, Director, Molecular and Applied Sciences Laboratory, Auburn University, 301 Wire Road, Office 260, Auburn, AL 36849, Andreas N. Kavazis, PhD, Professor, School of Kinesiology, Director, Muscle Biochemistry Laboratory, Auburn University, 301 Wire Road, Office 260, Auburn, AL 36849. indicates both authors equally contributed to the writing of this work (Breanna Mueller, Carlton D. Fox).

## Abstract

The effects of higher-load (HL) versus higher-volume (HV) resistance training (RT) on various molecular outcomes are similar. However, mitochondrial responses remain understudied. Therefore, the purpose of this study was to interrogate mitochondrial mRNA and protein responses to acute and chronic HL versus HV RT. Vastus lateralis biopsies from resistance trained males in two prior studies were assessed. In STUDY 1, 11 college-aged men completed an acute bout of either HL or HV RT exercises to failure. Biopsies were collected at PRE, 3 hours post-, and 6 hours post-exercise. In STUDY 2, 15 college-aged men participated in six weeks of supervised unilateral RT where each leg was assigned to either HL or HV RT. Biopsies were collected from both legs prior to and 72 hours following last training bout of the intervention. Biopsies from both studies were used to assess mitochondrial mRNAs, and STUDY 2 biopsies were assayed for mitochondrial proteins and CS activity. Results from both studies revealed several significant main effects of time but no significant interactions. Additionally, CS activity, a surrogate of mitochondrial content, decreased following chronic RT (p=0.016) but no interaction was observed between the HV and HL leg over time (p=0.882). In conclusion, while RT resulted in both acute mitochondrial mRNA as well as chronic CS activity and mitochondrial protein responses, there were no differences in the HL versus HV paradigms on these outcomes.

## Introduction

Skeletal muscle is a highly plastic tissue that responds to applied stimuli (Adams & Bamman, 2012; Bodine, 2013; Neufer et al., 2015). Perhaps the most well-studied of these adaptations is hypertrophy in response to mechanical overload (Roberts et al., 2023). Resistance training (RT) is the most accessible form of mechanical overload for humans. Thus, many studies have aimed at optimizing skeletal muscle hypertrophy via modifications to RT protocols. Two prominent modifications are the alteration of load (e.g. weight lifted) or volume (e.g. number of sets or repetitions performed) (Schoenfeld et al., 2017). As a result, studies examining training adaptations to high volume (HV) versus high load (HL) RT have gained widespread interest (Davies et al., 2022; Grisebach et al., 2024; Moir et al., 2011; Nitzsche et al., 2020). HL RT involves lifting heavier weight for fewer repetitions (e.g. 3-5 sets at 80% or more of one repetition maximum (1RM) for 3-5 repetitions) (Kraemer & Ratamess, 2004), while HV RT involves lifting lighter weight while performing more repetitions per set comparatively (e.g. sets containing >10 repetitions per set with < 65% of 1RM) (Krieger, 2010). Though HL training accomplishes greater increases in 1RM relative to HV training (Schoenfeld et al., 2015), both HL and HV training to failure results in similar hypertrophic adaptations (Mitchell et al., 2012; Morton et al., 2019).

Although strength and hypertrophy adaptations between HL and HV training have been compared (Burd et al., 2010; Mitchell et al., 2012; Morton et al., 2019; Vann et al., 2022), there is little evidence describing mitochondrial adaptations. Lim et al. (Lim et al., 2019) recruited college-aged males to perform 10-weeks of either 80FAIL training (80% 1RM to volitional fatigue), 30WM training (30% 1RM volume matched to 80FAIL), or 30FAIL training (30% 1RM to volitional fatigue) RT where PRE- and POST-intervention biopsies were examined for mitochondrial proteins indicative of mitochondrial volume, mitochondrial biogenesis, and remodeling. While Lim et al. reported a post-training increase in COX IV, a marker of mitochondrial capacity, there were no changes in markers of mitochondrial biogenesis (Lim et al., 2019). However, the authors reported HV training (30WM and 30FAIL) increased markers of mitochondrial remodeling (OPA1 and DRP1) compared to 80FAIL training. This finding is perhaps indicative of differential demands in energy production and mitochondrial activity between HL versus HV RT. Indeed, it has been suggested that HV RT might contribute to greater impacts to the mitochondria than HL RT (Parry et al., 2020). However, little research has been performed in this area.

Our group recently investigated transcriptional responses to a single bout of HL and HV training in previously trained college-age men (referred to as STUDY 1 herein) (Sexton et al., 2023). While these data were informative, genes related to mitochondrial biogenesis and remodeling were not investigated. Furthermore, using a six-week unilateral resistance training intervention in previously trained college-age men (referred to as STUDY 2 hereafter) we have shown elevation in integrative non-myofibrillar protein synthesis rates in the HV leg as compared to HL which may be indicative of changes in bioenergetic related proteins (Vann et al, 2022). Despite this, mitochondrial markers and CS activity were not assessed in the original investigation. Given that mitochondrial responses to HV or HL RT have scarcely been investigated, we leveraged specimens from the aforementioned studies to interrogate how HL and HV training affected markers of mitochondrial biogenesis and remodeling (Sexton et al., 2023; Vann et al., 2022). We hypothesized HV training would increase markers of mitochondrial biogenesis, mitochondrial remodeling, and CS activity relative to HL training.

## Methods

The methods outlined below are a partial report of relevant methods and procedures performed in STUDY 1 (Sexton et al., 2023) and STUDY 2 (Vann et al., 2022). A comprehensive description of the original data collections comprising this investigation can be found in the Sexton et al. (2023) and Vann et al. (2022) reports.

### Ethics approval and participant inclusion criteria

All procedures were approved by the Institutional Review Board at Auburn University (Protocols #20-081 MR 2003 and #19-245 MR 1907), and these studies conformed to the standards set forth by the latest revision of the Declaration of Helsinki except for not being registered as clinical trials. Participants were recruited verbally and through fliers posted at Auburn University. Eligible participants from both studies: i) self-reported ≥12 months of RT, ii) were free of cardiometabolic diseases (e.g., type II diabetes, severe hypertension), and iii) did not possess conditions precluding participation in the RT or the collection of skeletal muscle biopsies. Interested participants provided verbal and written informed consent to participate in the studies before the data collection procedures outlined below. Notably, 11 male participants were included in STUDY 1 analysis, and 15 separate participants were included in STUDY 2 analysis unless stated otherwise.

### Study designs and biochemical assays

Figure 1A displays a schematic of STUDY 1 (Sexton et al., 2023). In this study, 11 resistance-trained males made three visits to the laboratory. The first visit included a vastus lateralis muscle biopsy (PRE) and back squat and leg extension one repetition maximum (1 RM) test. These 1 RM values served to prescribe training loads for each of the following two visits during which participants performed four sets of back squats and leg extensions to failure using either 30% 1RM (HV) or 80% 1RM (HL) in a randomized, within-subject design. Following each acute bout of RT, vastus lateralis muscle biopsies were collected at 3- and 6-hrs post and all timepoints were processed for transcriptomic analysis.

**Figure 1.**
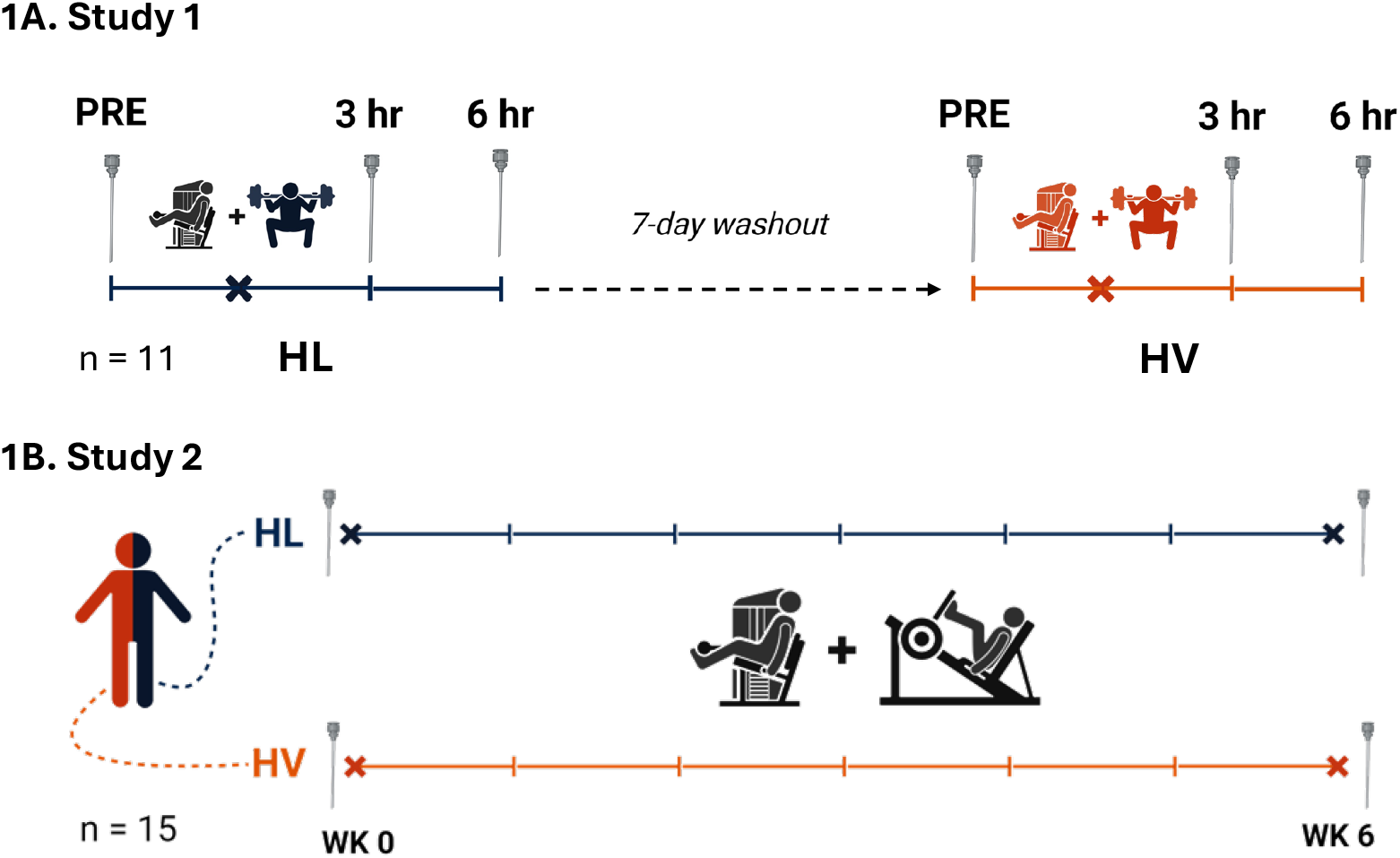
Study designs In study 1, participants performed a single bout of high volume (HV) resistance exercise (RE) or a single bout of high load (HL) RE in a randomized, crossover design with a 7-day washout period between testing sessions. Skeletal muscle biopsies were taken at pre, 3 hr and 6 hr after RE bout at each testing session. In study 2, participants performed 6 weeks of unilateral RT where the left and right legs of each participant were randomly assigned to perform either HV or HL resistance exercise. Skeletal muscle biopsies were taken at PRE and POST of the training intervention. Symbols and abbreviations: Needle symbol represents a biopsy; High volume (HV); High load (HL); Week (WK). Study design figures were created using BioRender (http://www.biorender.com).

### Data mining for mRNAs of interest from STUDY 1

Genes investigated in STUDY 1 were selected from a transcriptome-wide mRNA analysis that was previously attained and described by Sexton et al. (2023). To summarize, ∼10 mg of skeletal muscle was crushed in Trizol (VWR; Radnor, PA, United States) and centrifuged to isolate RNA. Next, purity was inspected using a NanoDrop Lite Spectrophotometer (Thermo Fisher, Waltham, MA, United States). Finally, samples were shipped on dry ice to a commercial laboratory for transcriptome-wide analysis using the Clariom S Assay Human mRNA array (North American Genomics; Decatur, GA, United States). Resultant .CEL files (raw data) were analyzed using the Transcriptome Analysis Console v4.0.2 (TAC; Thermo Scientific). Manual selection of mRNAs involved in mitochondrial biogenesis (PGC1α, TFAM, NRF1), fusion (MFN2, OPA1), fission (DRP1) and mitophagy (PINK1, PARKIN) were subjected to statistical analysis.

Figure 1B displays a schematic of STUDY 2 (Vann et al., 2022). Briefly, 15 resistance-trained males completed six weeks of total body RT (3x/week). Prior to the start of training, baseline vastus lateralis muscle biopsies were collected (PRE) and 1RM values for relevant RT exercises were determined. Lower body exercises included unilateral leg press and unilateral leg extension to allow for contrasting training schemes to be carried out within-subject. Left and right legs were randomly assigned to either HV or HL unilateral training. The HV leg began training performing 5 sets of 10 repetitions at 60% 1RM for each knee extensor exercise and increased training volume over 6 weeks ending at 10 sets of 10 repetitions at 60% 1RM at week 6. The HL leg began training performing 9 sets of 5 repetitions at 82.5% 1RM for each knee extensor exercise with training volume kept consistent while training load increased over 6 weeks, with week 6 training being performed at 95% 1RM. Vastus lateralis muscle biopsies were also collected 72 hours after the last bout (POST); both PRE and POST samples were processed for western blot analysis from 15 participants and qPCR analysis from 12 participants.

### Citrate synthesis activity assay

Muscle tissue stored in foils were removed from −80ºC and placed on a liquid nitrogen-cooled ceramic mortar. Crushed tissue (∼20 mg) was weighed on a laboratory scale exhibiting a sensitivity of 0.1 mg (Mettler-Toledo; Columbus, OH, USA), and tissue was quickly placed in 1.7 mL tubes containing 200 μL lysis buffer (25 mM Tris, pH 7.2, 0.5% Triton X-100, 1x protease inhibitors). Samples were homogenized using tight-fitting hard-plastic microtube pestles on ice and centrifuged at 1,500 g for 10 minutes at 4°C. Supernatants were collected and placed in new 1.7 mL microtubes on ice and subsequently assayed for protein concentration using a commercially available BCA kit (Thermo Fisher Scientific; Waltham, MA, USA). Samples were assayed in duplicate using a microplate assay protocol (20 μL of 5x diluted sample + 200 μL Reagent A + B). Samples were then stored at −80ºC until assays described below.

Tissue lysates obtained above were batch processed for citrate synthase (CS) activity as previously described by our laboratory (Kephart et al., 2015). This metric was used as a surrogate for mitochondrial content per the findings of Larsen et al. (Larsen et al., 2012), suggesting CS activity correlates with transmission electron micrograph images of mitochondrial content (r = 0.84, p < 0.001). The assay principle is based upon the reduction of 5,50-dithiobis (2-nitrobenzoic acid) (DTNB) at 412 nm (extinction coefficient 13.6 mmol/L/cm) coupled to the reduction of acetyl-CoA by the CS reaction in the presence of oxaloacetate. Skeletal muscle protein (5 µg) from each biopsy was assayed in duplicate where samples were added to a mixture composed of 0.125 mol/L Tris–HCl (pH 8.0), 0.03 mmol/L acetyl-CoA, and 0.1 mmol/L DTNB. The reaction was initiated by adding 5 μL of 50 mmol/L oxaloacetate and the absorbance change was recorded for 1 min. The coefficient of variation for all duplicates was 6.1%.

### Western blotting

Protein expression data obtained from STUDY 2 was attained via Western blot analysis as previously detailed by Vann et al. (Vann et al., 2021). Briefly, lysates obtained were prepared for Western blotting using 4x Laemmli buffer at one μg/μL. Following sample preparation, 15 μL samples were loaded onto pre-casted gradient (4–15%) SDS-polyacrylamide gels (Bio-Rad Laboratories, Hercules, CA, USA) and subjected to electrophoresis (180 V for 45– 60 min) using pre-made 1x SDS-PAGE running buffer (VWR International, Randor, PA, USA). Proteins were subsequently transferred (200 mA for 2 h) to polyvinylidene difluoride membranes (PVDF) (Bio-Rad Laboratories, Hercules, CA, USA), Ponceau stained, and imaged to ensure equal protein loading between lanes. Membranes were then blocked for 1 h at room temperature with 5% nonfat milk powder in Tris-buffered saline with 0.1% Tween-20 (VWR International). Rabbit anti-peroxisome proliferator-activated receptor gamma coactivator 1-alpha (PGC1α, 1:1000; catalog #: GTX37356; GeneTex, Irvine, CA, USA), rabbit anti-nuclear respiratory factor 1 (NRF1, 1:2000, catalog# GTX103179, 11168915, GeneTex), rabbit anti-transcription factor A, mitochondria (TFAM, 1:2000, catalog# H00007019-D01P,171 5621, Abnova Corporation, Taoyuan City, Taiwan), rabbit anti-mitofusin 2, (MFN2, 1:2000, catalog# 3882-100; BioVision, Milpitas, CA), rabbit anti-dynamin-related protein 1 (DRP1, 1:2000, catalog# NB110-55288SS; Novus Biologicals USA, Littleton, CO), rabbit anti-PRKN parkin RBR E3 ubiquitin protein ligase (PARKIN, 1:2000, # 2132, 10693040, Cell Signaling, Danvers, CO, USA), rabbit anti-PTEN-induced kinase 1 (PINK1, 1:2000, catalog# 6946, 11179069, Cell Signaling) were incubated with membranes overnight at 4 °C in TBST with 5% bovine serum albumin (BSA). The following day, membranes were incubated with horseradish peroxidase (HRP)-conjugated anti-rabbit IgG (catalog #: 7074; Cell Signaling) in TBST with 5% BSA at room temperature for 1 h (secondary antibodies diluted 1:2000). Membrane development was performed using an enhanced chemiluminescent reagent (Luminata Forte HRP substrate; EMD Millipore, Billerica, MA, USA), and band densitometry was performed using the ChemiDoc Touch gel documentation system and associated software (Bio-Rad Laboratories, Hercules, CA, USA). Raw densitometry values for each target were divided by whole-lane Ponceau densities, and these data were statistically analyzed between groups.

### Quantitative polymerase chain reaction

The mRNA expression from 12 samples from STUDY 2 were attained via qPCR analysis as previously described by Romero et al (2018). In short, isolated RNA was reverse transcribed into cDNA for quantitative polymerase chain reaction (qPCR) analysis using 2 mg of RNA with cDNA synthesis reagents (Quanta Biosciences, Gaithersburg, MD) in accordance with the manufacturer’s recommendations. Quantitative PCR was performed using SYBR green chemistry (Quanta Biosciences). Calculations for qPCR were also performed as previously described in Romero et al. (Romero et al., 2018).

### Statistical analysis

Data collected from both studies was statistically analyzed using GraphPad Prism v10.2.2 (San Diego, CA, United States) (https://www.graphpad.com/). Normality was assessed using Shapiro Wilks and nonnormal data (TFAM protein were log transformed). Two-way, repeated measures ANOVAs were used to determine the interaction between and/or main effects of time (STUDY 1: PRE, 3 hr and 6 hr; STUDY 2: PRE and POST) and condition (STUDY 1: HL vs HV; STUDY 2 and HL vs HV). Tukey’s post-hoc pairwise comparisons were performed when interactions reached statistical significance to further investigate the simple main effects of time and/or condition.

## Results

### STUDY 1: Effects of acute bouts of HV versus HL on mRNA markers of mitochondrial biogenesis and remodeling

All markers of mitochondrial biogenesis demonstrated significant main effects of time, but no condition*time interactions were observed. Specifically, PGC1α mRNA increased from PRE to 3 hr (p < 0.001) and 6 hr (p ≤ 0.001) (Fig. 2a), NRF1 mRNA expression decreasing from PRE to 3 hr but returning to baseline at 6 hr (p = 0.021, Fig. 2b), and TFAM mRNA expression increasing from PRE to 6 hr (p = 0.035, Fig. 2c).

**Figure 2.**
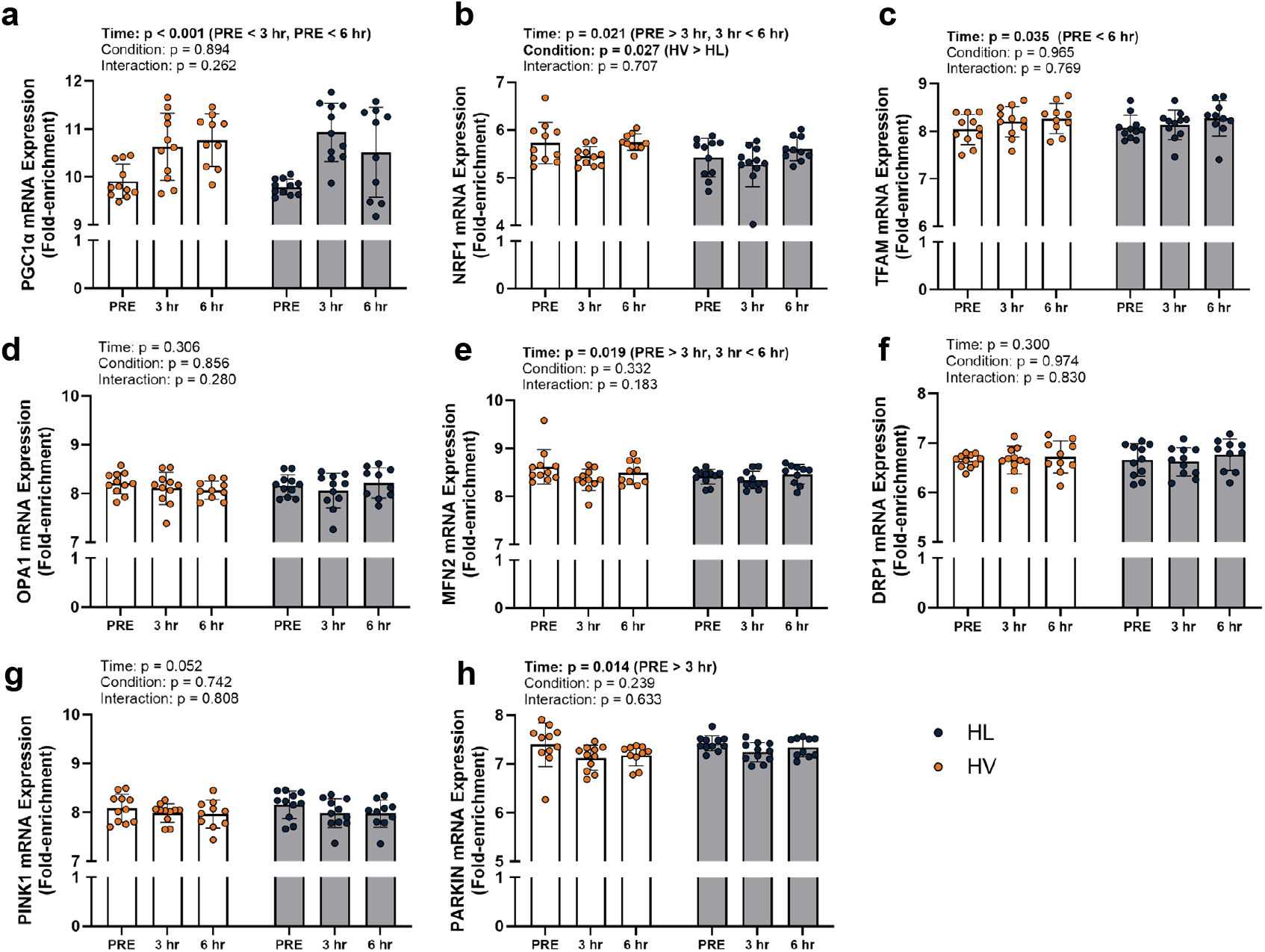
STUDY 1 mRNA expression data Figure 2 depicts mRNA expression from 11 participants (n = 11) reported by fold-enrichment. Orange data points and dark gray bars represent the high-volume resistance training while dark blue data points and light gray bars represent high load resistance training. Each condition includes PRE, 3 hours POST and 6 hours POST time points. Abbreviations: peroxisome proliferator-activated receptor gamma coactivator-1 alpha (PGC1α), mitochondrial transcription factor A (TFAM), nuclear respiratory factor 1 (NRF1), mitofusion 2 (MFN2), optic atrophy 1 (OPA1), dynamin-related protein 1 (DRP1), PTEN induced putative kinase 1 (PINK1), high volume (HV), high load (HL).

Among all markers of mitochondrial remodeling, only two markers demonstrated main effects of time with no main effects of RT condition or condition*time interactions. MFN2 mRNA expression decreased from PRE to 3 hr but returned to baseline by 6 hr (p = 0.019, Fig. 2e) and PARKIN mRNA expression decreased from PRE to 3 hr (p= 0.014, Fig. 2h).

### STUDY 2: A summary of general training characteristics and adaptations to chronic RT training

A full report of general training characteristics and adaptations were previously reported by (Vann et al., 2022), but these details are provided here for convenience. In summary, a significant interaction was evident for vastus lateralis muscle cross-sectional area (assessed via magnetic resonance imaging; interaction p=0.046), where HV increased this metric from PRE to POST (+3.2%, p=0.018), but HL training did not (−0.1%, p=0.475). Interestingly, six-week integrated non-myofibrillar protein synthesis rates were also higher in the HV versus HL leg (p=0.018), while no difference between legs existed for integrated myofibrillar protein synthesis rates (p=0.687).

### STUDY 2: Effect of chronic HV versus HL RT on mRNA expression on markers of mitochondrial biogenesis and remodeling

Six weeks of chronic RT did not affect mRNA expression of genes related to mitochondrial biogenesis and remodeling (p ≥ 0.297) (Figure 3).

**Figure 3.**
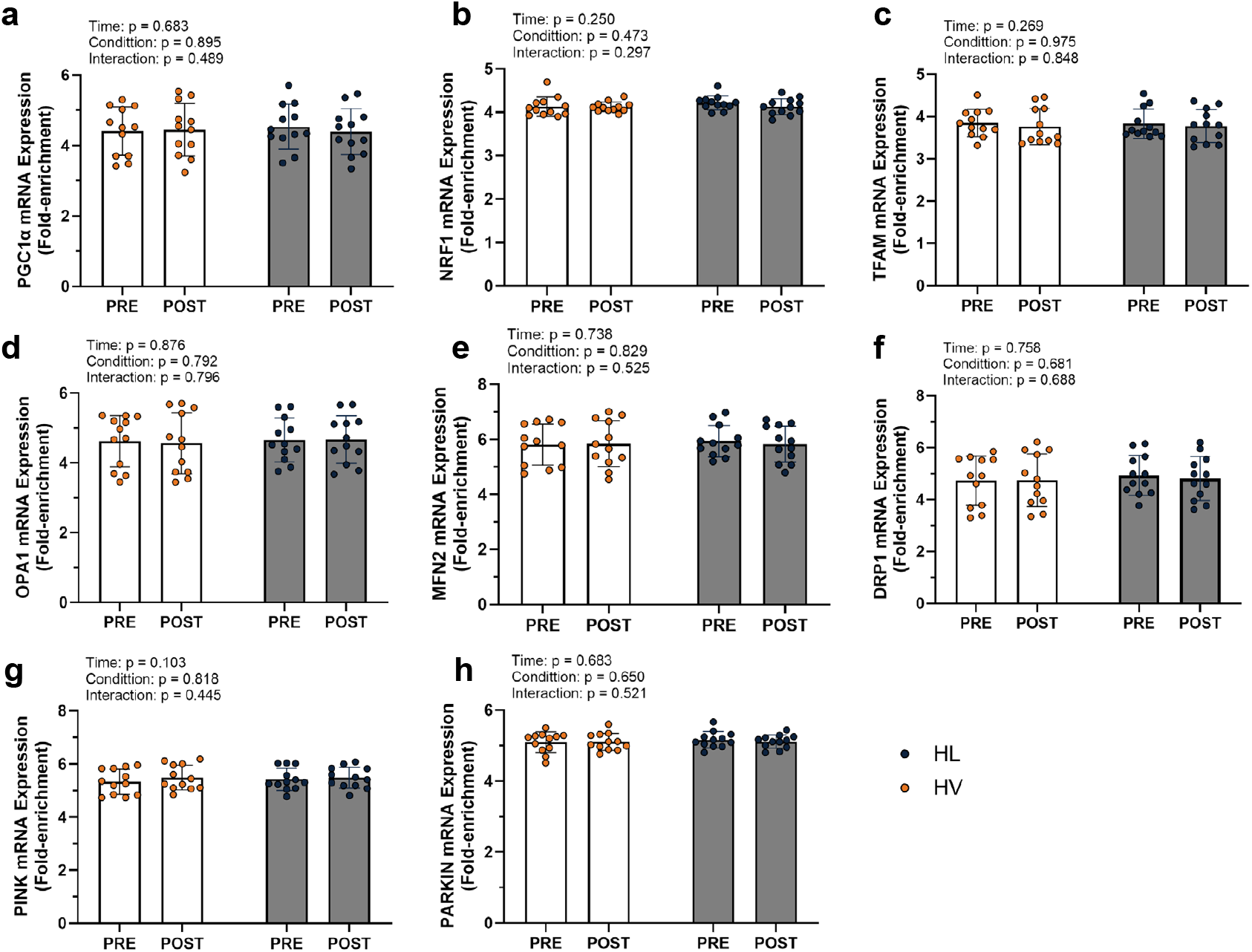
STUDY 2 mRNA expression data Figure 3 depicts mRNA expression from 12 participants (n = 12) reported by fold-enrichment. Orange data points and dark gray bars represent the high-volume resistance training while dark blue data points and light gray bars represent high load resistance training. Each condition includes PRE and POST time points. Abbreviations: peroxisome proliferator-activated receptor gamma coactivator-1 alpha (PGC1α), nuclear respiratory factor 1 (NRF1), mitochondrial transcription factor A (TFAM), optic atrophy 1 (OPA1), mitofusion 2 (MFN2), dynamin-related protein 1 (DRP1), PTEN induced putative kinase 1 (PINK1), high volume (HV), high load (HL).

### STUDY 2: Effects of chronic HV versus HL RT on mitochondrial markers

Interrogation of proteins associated with mitochondrial biogenesis in response to chronic HV and HL RT revealed only main effects of time where PGC1α (p < 0.001, Fig. 4a) and TFAM (p = 0.032, Fig. 4c) protein expression decreased from PRE to POST while NRF1 protein expression increased from PRE to POST (p = 0.001, Fig. 4b). However, there were no significant condition*time interactions observed.

**Figure 4.**
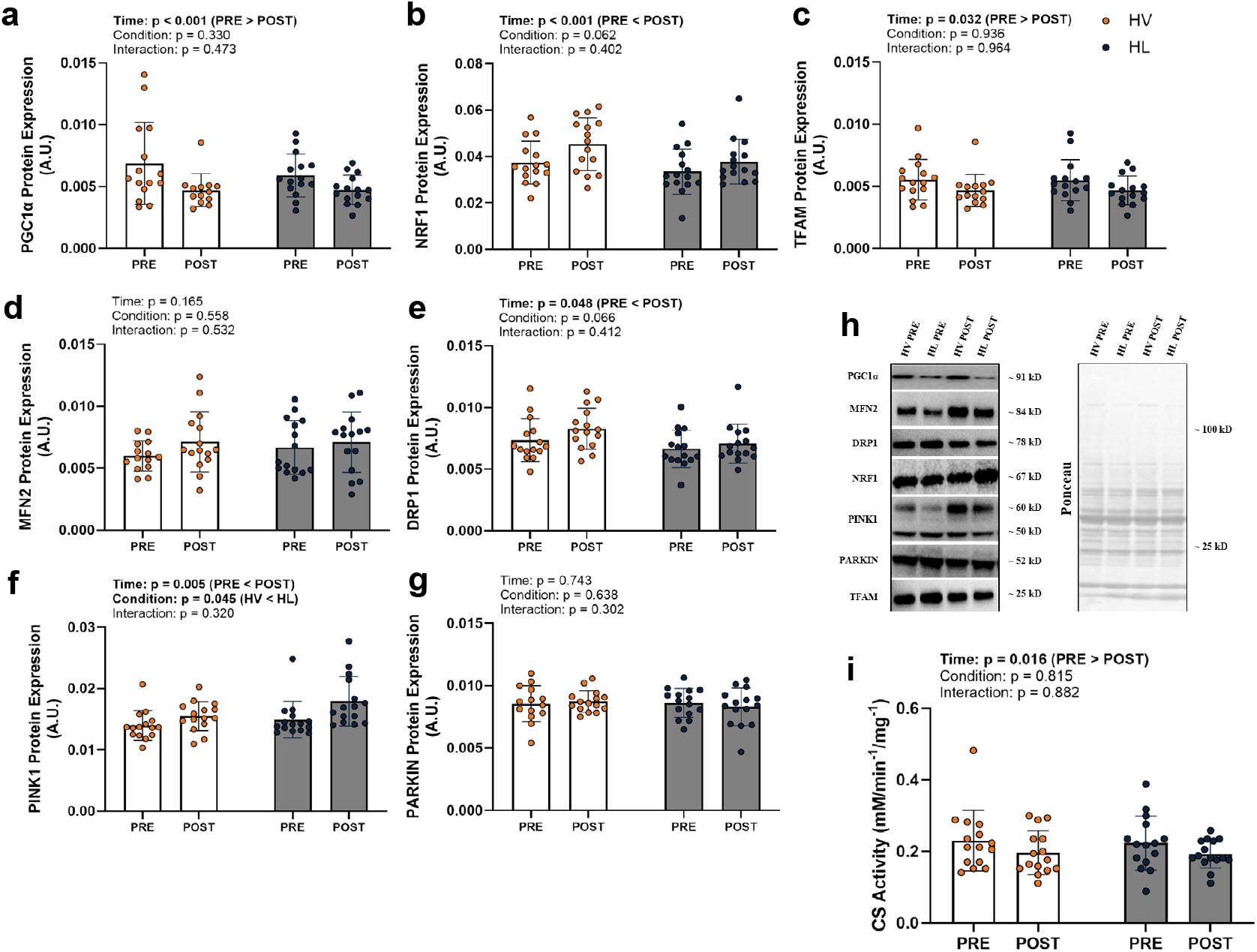
STUDY 2 Protein expression and CS activity data Figure 4 depicts protein levels and citrate synthase activity from 15 participants (n = 15). Orange data points and dark gray bars represent the high-volume resistance training while dark blue data points and light gray bars represent high load resistance training. Each condition includes PRE and POST time points. Each protein marker is accompanied by a representative western blot image located to the right of each graph. Abbreviations: peroxisome proliferator-activated receptor gamma coactivator-1 alpha (PGC1α), mitochondrial transcription factor A (TFAM), nuclear respiratory factor 1 (NRF1), mitofusion 2 (MFN2), optic atrophy 1 (OPA1), dynamin-related protein 1 (DRP1), PTEN induced putative kinase 1 (PINK1), arbitrary units (A.U.), kilodalton (kD), citrate synthase activity (CS Activity), high volume (HV), high load (HL).

Among markers of mitochondrial remodeling in response to chronic bouts of HV and HL RT, DRP1 and PINK1 demonstrated main effects of time with protein expression increased at POST for both markers (p = 0.005 and p = 0.048, respectively, Fig. 4e/f). However, there were no significant condition*time interactions observed.

CS activity in response to chronic bouts of HV and HL RT demonstrated a main effect of time where CS activity decreased from PRE to POST (p = 0.016, Fig. 4i). However, no significant condition*time interactions observed.

## Discussion

The purpose of this study was to investigate the acute transcriptional and chronic translational responses of mitochondrial specific genes/proteins to HL versus HV RT. Specifically, we accomplished this by interrogating previously collected samples from two groups of participants completing either acute or chronic HV and HL RT. Based on limited data, we hypothesized that HV training would have a more robust effect on increasing mitochondrial markers than HL training. However, our findings herein do not support this hypothesis.

Our primary finding is that acute RT transiently increases transcriptional markers of mitochondrial biogenesis and remodeling. We posit these data add further support that RT provides some benefit to mitochondrial adaptations in addition to more characterized benefits such as increased lean mass and strength (Di Leo et al., 2023; Roberts et al., 2023; Taivassalo & Haller, 2005; Zaenker et al., 2018). A similar finding was reported by Burd and colleagues (Burd et al., 2012) where mitochondrial protein fractional synthesis rates increased at 6- and 24-hours post-RT bout and PGC1α mRNA expression increased 6 hours post-RT bout. However, it should be noted that the effect of chronic RT on mitochondrial markers is mixed (Groennebaek & Vissing, 2017; Kon et al., 2014; Porter et al., 2015; Tang et al., 2006). Specifically, our observation of a decrease in CS activity following resistance training suggests that mitochondrial volume density decreased. It is important to emphasize that, as muscle hypertrophy ensues with resistance training, no change in tissue CS activity levels likely indicates that mitochondrial volume density proportionally increases with cell growth. Conversely, if muscle CS activity levels decrease with resistance training, this may reflect a lag in the expansion of mitochondrial volume density relative to cell growth (Parry et al., 2020). In this regard, a comprehensive review by Groennebaek et al. cites numerous resistance training studies reporting no change or a decrease in muscle CS activity, and we have similarly shown that CS activity levels decrease in trained and untrained participants following periods of resistance training (Groennebaek & Vissing, 2017; Haun et al., 2019; Roberts et al., 2018). These collective reports have led us to speculate that changes in muscle CS activity with resistance training are likely dependent upon the training status and age of the participant, where increases in this marker are typically observed in untrained subjects (especially older untrained participants) (Parry et al., 2020). Recent data from our laboratory, however, adds insight to this area. We assessed CS activity changes following 10 weeks of resistance training in untrained participants and compared these data to what was observed via histology using a mitochondrial specific antibody (TOMM20) (Ruple et al., 2021). Our histological analysis indicated mitochondrial volume density significantly increased, and in fact, outpaced muscle fiber growth, whereas tissue CS activity levels were not altered. Moreover, no association was evident when examining the pre-to-post intervention changes in both metrics. Given these data, CS activity assay may lack the sensitivity to track changes in mitochondrial volume density with RT interventions, especially in studies involving previously trained participants where adaptations are likely less robust. In line with this hypothesis, the current data suggest that mitochondrial volume density is marginally affected with either form of training, and this may have been due to the training status of the participants.

Our observations also suggested that the mRNA response to an acute bout of resistance exercise is not always indicative of translational response to chronic resistance exercise. For instance, though PGC1α mRNA expression was elevated at 3 hr and 6 hr following an acute bout of HV or HL RT, chronic RT did not result in a complimentary increase in PGC1α protein expression. In fact, PGC1α protein expression was decreased following chronic RT. PARKIN and MFN2 also demonstrated changes in mRNA expression following an acute bout of resistance exercise. However, neither gene demonstrated any change in protein expression following chronic resistance exercise. Conversely, neither DRP1 nor PINK1 showed any transcriptional response to an acute bout of resistance exercise, though chronic RT resulted in statistically significant changes in protein expression for these genes. While these data suggest a disconnect between the transcriptional response to acute training versus the translational response with chronic training it is critical to note that these data were collected from different participants undergoing different training paradigms and thus are difficult to compare. Participant and protocol differences notwithstanding, these findings and the argument presented by Miller and colleagues reiterate the notion that mRNA expression alone is not representative of longer-term adaptations (Miller et al., 2016). While this may appear to be yet another challenge in the accurate determination of molecular signaling events following RT, this disconnect may shed light on key mechanisms of molecular modulation in RT adaptation. Considering the divergence observed between the initial transcriptional response and the subsequent translational response of specific genes, we postulate that additional factors affecting the molecular response to resistance exercise may be at play. It is possible that epigenetic modifications (e.g. micro RNAs, long non-coding RNAs, DNA methylation, histone modifications) play a role in the protein expression of such transcribed mitochondrial mRNAs (Mcleod et al., 2024; Stokes et al., 2020). Research within this field of study has already demonstrated an effect of epigenetic modification in response to RT (Seaborne & Sharples, 2020; Sexton et al., 2023). Given these discordant findings and the established RT responsiveness of epigenetic factors, investigations into the epigenetic modifications that might impact the protein expression (or lack thereof) of transcribed mitochondrial mRNAs in response to RT are warranted.

## Limitations

One limitation was our participant population. Considering that only young, trained, male participants were utilized in this investigation, these results cannot be generalized to female or untrained populations. Another limitation of this study was the short training duration, which might not have provided sufficient stimulus to observe differences between each training condition in previously trained participants. Likewise, although the HV leg in STUDY 2 did experience significantly more volume (Vann et al., 2022), this was only an 11% difference. This lack of appreciable training volume differences between legs may have been a chief reason why we observed several null findings. In this regard, longer training studies that measure these markers with more pronounced volume differences between legs are needed to determine how the muscle-metabolic milieu adapts in response to different RT paradigms. Additionally, it is critical to note that while HV versus HL protocols were implemented in a similar cohort of participants, these data were collected from separate studies with individuals performing differential training protocols. Therefore, it is difficult to make exact comparisons for mRNA and protein expression data.

## Conclusions

Although RT resulted in both acute transcriptional (mRNA) and chronic CS activity and translational (protein) responses, there was little effect of the RT paradigm (HV or HL) in markers of mitochondrial biogenesis and remodeling. These results indicate that RT can alter mitochondrial markers both acutely and chronically and suggest that HV and/or HL RT can both protocols be leveraged to promote mitochondrial adaptations. Additionally, we observed that chronic protein level responses did not necessarily reflect transcriptional responses, though these data were collected in a separate cohort of participants. Nonetheless, these results point to possible epigenetic modifications to the molecular events following acute and chronic resistance exercise. Therefore, additional measures besides mRNA and protein expression must be taken to comprehensively assess molecular events following resistance exercise.

## CONFLICTS OF INTEREST

None of the authors have conflicts of interest in relation to the current data. In the interest of full transparency, the laboratory of MDR does perform contracted studies from industry various sponsors. MDR has also served as a paid consultant to various industry entities in line with Auburn University’s Conflict of Interest Policies. SMP has received grant funding from the Canadian Institutes of Health Research, the National Science and Engineering Research Council of Canada, the US National Institutes of Health, Roquette Frères, Nestlé Health Sciences, FrieslandCampina, the US National Dairy Council, Dairy Farmers of Canada, Myos, and Cargill. SMP has received travel expenses and honoraria for speaking from Nestle Health Sciences, Optimum Nutrition, Nutricia, and Danone. SMP holds patents licensed to Exerkine Inc. but reports no financial gains from these patents or otherwise.

## ACKNOWLEDGEMENTS

The authors would like to thank the participants for their dedication to executing this study. We would also like to thank Dr. Shelby Osburn, Dr. Morgan Smith, Dr. Melissa Rumbley, Brian Ferguson, Samantha Slaughter, Andy Cao, Max Coleman, Max Michel, Megan Edwards, and Sullivan Clement for their assistance in collecting data.

## FUNDING

Funding for assays and participant compensation was provided through discretionary laboratory funds from M.D.R. and A.N.K.

